# An effect size distribution analysis of heart rate variability studies: Recommendations for reporting the magnitude of group differences

**DOI:** 10.1101/072660

**Authors:** Daniel S. Quintana

## Abstract

The calculation of heart rate variability (HRV) is a popular tool used to investigate differences in cardiac autonomic control between population samples. When interpreting effect sizes to quantify the magnitude of group differences, researchers typically use Cohen's guidelines of small (0.2), medium (0.5), and large (0.8) effects. However, these guidelines were only proposed for use when the effect size distribution (ESD) was unknown. Despite the availability of effect sizes from hundreds of HRV studies, researchers still largely rely on Cohen's guidelines to interpret effect sizes. This article describes an ESD analysis of 297 HRV effect sizes from case-control studies, revealing that the 25th, 50th, and 75th effect size percentiles correspond with effect sizes of 0.25, 0.5, and 0.84, respectively. The ESD for separate clinical groups are also presented. The data suggests that Cohen's guidelines underestimate the magnitude of small and large effect sizes for the body of HRV case-control research. Therefore, to better reflect observed HRV effect sizes, the data suggest that effect sizes of 0.25, 0.5, and 0.85 should be interpreted as small, medium, and large effects. Researchers are encouraged to use the ESD dataset or their own collected datasets in tandem with the provided analysis script to perform bespoke ESD analyses relevant to their specific research area.

## 1. Introduction

The calculation of heart rate variability (HRV) – the complex modification of the heart rate over time – has become a widely adopted tool to non-invasively index autonomic control of the heart rate (Billman, 2011). The comparison of HRV between a population of interest and control group is a popular approach used to assess differences in cardiac autonomic control. The identification of group differences in HRV has been used to better understand disease aetiology, psychological phenomena, and the increased risk of frequently comorbid illnesses, such as cardiovascular disease (Kemp and Quintana, 2013). Indeed, several meta-analyses have reported reduced HRV in a range of clinical populations in comparison to control groups (e.g., Alvares et al., 2016; Koenig et al., 2016; McIntosh, 2016).

When reporting the comparison of an independent variable between groups, many researchers place a disproportionately large emphasis on statistical significance, despite the weaknesses of this approach (Cumming, 2013). For instance, *p*-values do not give an accurate summary of the magnitude of observed differences between groups, only whether an effect exists. Moreover, *p*-values cannot be used to compare effects between studies that have different sample sizes. Finally, it can difficult to understand group difference magnitude when an unfamiliar variable is reported – a common occurrence in HRV research given that there are over 70 different published metrics (Bravi et al., 2011; Smith et al., 2013).

Given these issues, the American Psychological Association has recommended the reporting of effect sizes in manuscripts, highlighting that this practice is “essential to good research” (Wilkinson, 1999, p. 599). Unlike *p*-values, effect sizes communicate the magnitude of group differences, which can be compared between studies using common metrics. When reporting effect sizes, researchers almost always use Cohen’s conventions when interpreting effect magnitude - effect sizes of Cohen’s *d* (or the closely related Hedges’ *g* statistic) equal to 0.2, 0.5, and 0.8 are considered small, medium, and large effects, respectively. These thresholds were derived from Cohen’s suggestion that medium effects represent differences that are likely to be visible to the naked eye, which happens to approximate the average effect across fields, and large and small effects should be equidistant from medium effects (Cohen, 1992). Using guidelines designed to represent the average across fields may underestimate or overestimate the true distribution of effects in a specific research field, which is likely to have its own unique effect size distribution (ESD).

Despite the wide use of Cohen’s guidelines within the biobehavioral sciences, research has yet to establish the distribution of effect sizes in HRV case-control research. While effect size distributions have been investigated in other areas of psychology (Gignac and Szodorai, 2016), this is the first article to describe a systematic ESD analysis for a widely used biobehavioral variable. Moreover, this article also provides a companion script to perform all aspects of an ESD analysis using R. By calculating empirically derived effect size distributions, HRV researchers can better understand and communicate the magnitude of their effects guided by prior research in the field.

## 2. Methods

Effect sizes were extracted from meta-analyses that reported the synthesis of HRV case control studies comparing disorder populations with control groups. Meta-analyses were identified using the following search string in the PubMed database entered August 27, 2016: ((“meta analytic”[Title] OR “meta analysis”[Title]) AND (“heart rate variability”[Title] OR “autonomic”[Title] OR “parasympathetic”[Title] OR “vagal”[Title])). This search yielded 44 articles. Each article was examined for eligibility, which left 17 meta-analyses from which samples sizes were extracted (Supplementary Figure 1; Supplementary Table 1). Hedges’ *g* values for each study included in eligible meta-analyses were extracted, along with group sizes. When meta-analyses reported Cohen’s *d*(n = 2), these values were converted to Hedges’ *g* for consistency. Any negative Hedges’ *g* values were entered as positive values as the interest of this study was to determine the distribution of group difference effects, rather than the direction of effects. Hierarchical inclusion criteria were implemented to prevent the duplication of effect-size estimates: when different meta-analyses reported effect sizes from the same studies, effect sizes from the meta-analyses with the smaller amount of effect sizes were discarded. When multiple HRV measures were reported, the measure with the largest number of effect sizes was extracted. If these numbers were equivalent, the root mean square successive differences (RMSSD) value was chosen, as this measure is less susceptible to confounding respiratory effects (Penttilä et al., 2001; Quintana et al., 2016). Two unrealistically large effect sizes (Hedges’ *g*> 3) were removed from the sample.

To examine the distribution of effect sizes, a range of percentiles were calculated for all HRV studies and HRV study subgroups. The percentiles of particular interest are the 50^th^, as this represents the average effect size, and the 25^th^ and 75^th^ percentiles, as these are equidistant from the average effect size (Cohen, 1992). Consistent with Cohen’s (1992) recommendation to visually inspect of effect size differences to help determine effect magnitudes, a series of histograms were constructed to visualise differences between two hypothetical populations associated with the derived small, medium, and large effect sizes.

A one-sided contour-enhanced funnel plot, in which the effect size is plotted against standard error with added contours representing key levels of statistical significance (*p* = 0.1, *p* = 0.05, *p* = 0.01; Peters et al., 2008), was also constructed to examine if statistically significant effects are overrepresented in HRV research, which could bias the ESD. A majority of studies lying within “significance contours” may be indicative of inflation bias – popularly known as “*p*-hacking” – whereby investigators apply various analytical approaches until reaching a statistically significant result (Simmons et al., 2011). The dataset and script to perform all described analyses using the R statistical package is available at http://osf.io/rwj4h.

## 3. Results

A total of 297 effect sizes were extracted. The 25^th^ (small effect), 50^th^ (medium effect), and 75^th^ (large effect) percentiles corresponded to Hedges’ *g* values of 0.25, 0.5, and 0.84 (Fig. 1). In other words, a Hedges’ *g* of 0.5 is the average effect size across HRV case-control studies, consistent with Cohen’s guidelines. Moreover, this effect size is clearly visible to the naked eye (Fig. 2), fulfilling Cohen’s original criterion for a medium effect size. However, Cohen’s guidelines for small and large effect sizes appear to underestimate the empirically derived thresholds by 0.05.

**Figure 1.**
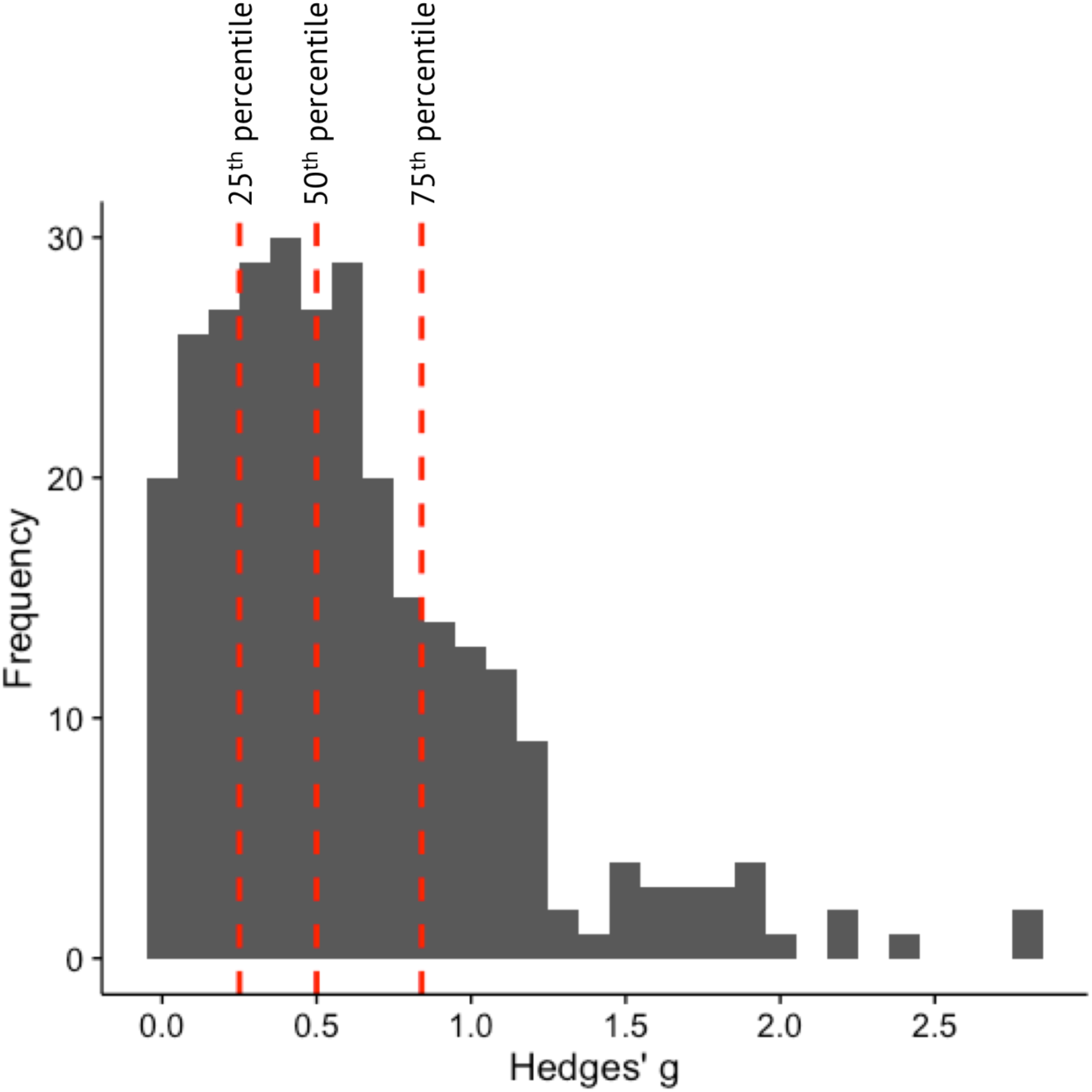
The distribution of 297 effect sizes from HRV case-control studies. The 25^th^, 50^th^, and 75^th^ percentiles (dashed red lines) represent the calculated thresholds for small (0.25), medium (0.5), and large (0.84) effects.

**Figure 2.**
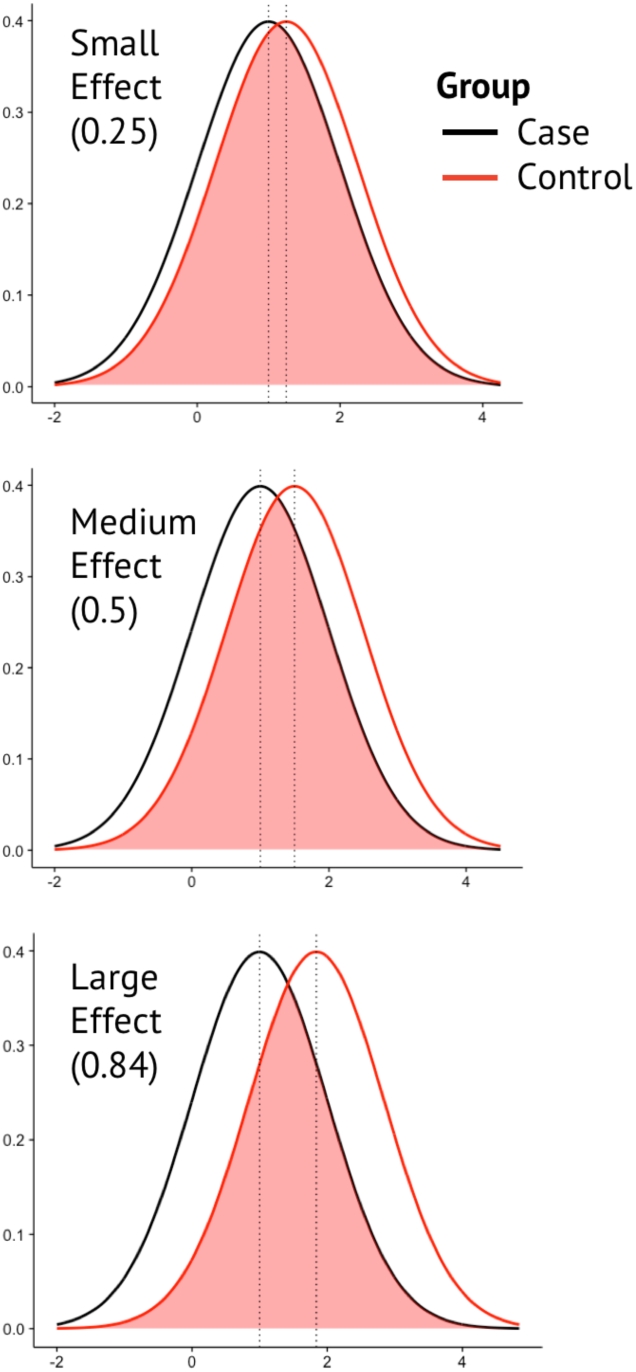
Histograms depicting differences between two hypothetical populations associated with the small, medium and large effect sizes derived from the present analysis. According to Cohen (1992), a medium effect should be clearly visible to the naked eye.

The small, medium, and large effect percentiles and effect size distributions for HRV study subgroups are presented in Table 1 and Figure 3 (see Supplementary Table 2 for a larger range of percentiles). Notable deviations from Cohen’s guidelines include the reduced large effect size percentile for anxiety disorders (0.63) and all effect size percentiles for psychosis spectrum disorders, which are much larger than commonly cited guidelines (small effect = 0.56, medium effect = 0.78, large effect = 1.1; Fig. 3B). Finally, visual inspection of the contour-enhanced funnel plot did not reveal an obvious over-representation of studies lying in the “significance contours” (Fig. 4). This is consistent with a low likelihood of discipline-wide inflation bias.

**Figure 3.**
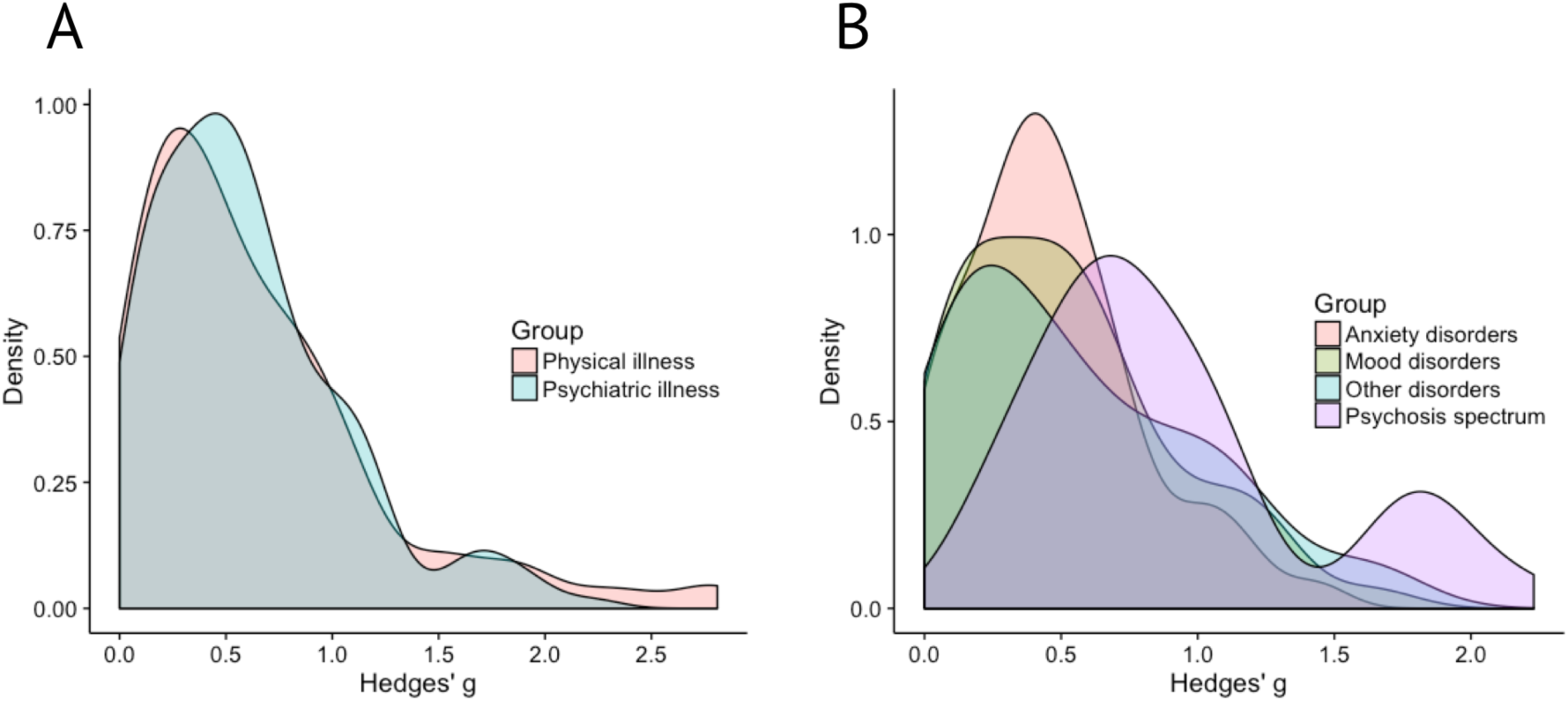
Density plots illustrating the distribution of Hedges’ *g*. The distributions of effect sizes from studies assessing physical and psychiatric groups are roughly equivalent (A). However, the distributions of effect sizes from psychiatric disorder groupings illustrate the larger average effect size of studies investigating psychosis spectrum disorders (B).

**Figure 4.**
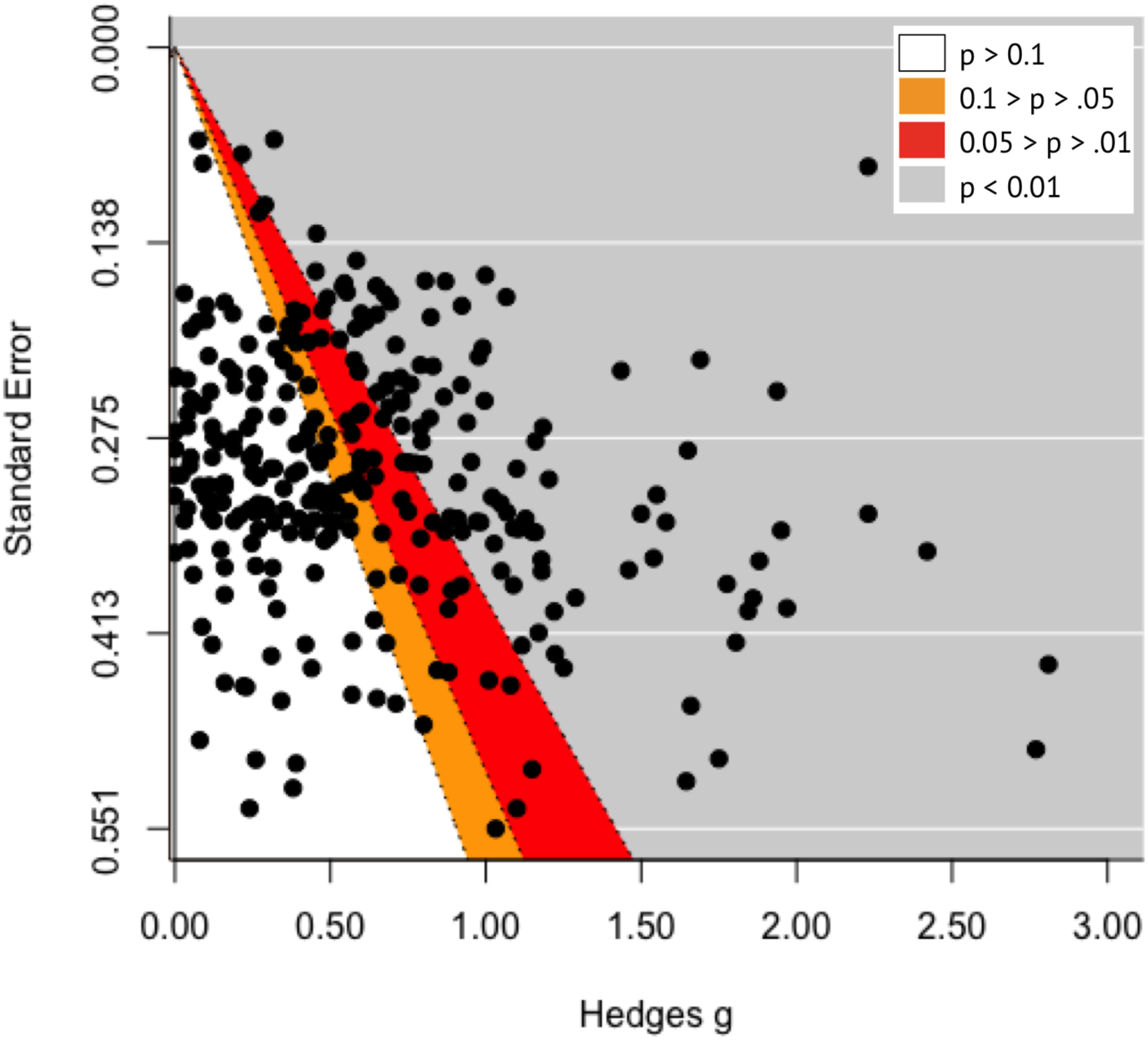
A one-sided contour-enhanced funnel plot, whereby effect sizes are plotted against standard errors, was constructed to explore if statistically significant effects are overrepresented in HRV research. Statistical significance can be calculated for a combination of effect size and standard error (assuming all studies used two-sided tests with a significance criterion of α = 0.05). Thus, contours representing key levels of statistical significance can be superimposed on the plot. As there was no pronounced over-representation of studies lying within the orange and red significance contours this suggests a low likelihood of discipline-wide inflation bias.

**Table 1.** Effect size percentiles.

|  | 25% | 50% | 75% |
| --- | --- | --- | --- |
| Cohen's suggested guidelines | 0.2 | 0.5 | 0.8 |
| HRV - all studies (k = 297) | 0.25 | 0.5 | 0.84 |
| HRV - psychiatric illnesses (k = 187) | 0.26 | 0.51 | 0.81 |
| HRV - physical illnesses (k = 110) | 0.25 | 0.49 | 0.88 |
| HRV - Anxiety disorders (k = 55) | 0.25 | 0.43 | 0.63 |
| HRV - Mood disorders (k = 53) | 0.19 | 0.45 | 0.69 |
| HRV - Psychosis spectrum disorders (k = 44) | 0.56 | 0.78 | 1.1 |
| HRV - Chronic pain (k = 73) | 0.19 | 0.47 | 0.8 |

## 4. Discussion

When Cohen first suggested effect size guidelines, he intended these to be used only when effect size distributions were unavailable (Cohen, 1988). Despite the availability of hundreds of HRV studies, many researchers – including the present author (e.g., Chalmers et al., 2016) – still rely on Cohen’s 1988 guidelines to interpret effect sizes. Consequently, the goal of the present research was to assess the ESD of HRV case-control studies to determine whether empirically derived interpretation guidelines better represent HRV studies. The data suggests that Cohen’s suggested guidelines might slightly underestimate the magnitude of small and large effect sizes in HRV case-control research, which is especially apparent for HRV research in psychosis spectrum disorders populations.

There are some limitations to the present investigation worth briefly noting. First, while extracting effect sizes from published meta-analyses provides an efficient means of collecting a large quantity of effect size data, this approach would have missed HRV studies not included in the 17 eligible meta-analyses. However, the collection of 398 effect sizes is likely representative of the overall distribution of effect sizes. Second, the comparison of HRV effect sizes from intervention studies was beyond the scope of the paper, so it not known if the ESD of these studies differ from the case-control ESD.

In summary, future research investigating case-control differences in HRV would benefit from using these empirically derived effect size distributions to convey group difference magnitudes. In view of the data, it is recommended that effect sizes of 0.25, 0.5, and 0.85 should be used as thresholds to interpret small, medium, and large effects if the researchers wish to use thresholds applicable to a representative sample of HRV studies across populations. Alternatively, researchers can perform their own ESD analysis on subsets of the data that are more relevant for their research with the provided analysis script and dataset, or create their own datasets for analysis using the methods described herein. Finally, ESD analyses can also facilitate more accurate power analyses when planning future research, as the effect sizes entered into these analyses are likely to better represent HRV research.

